# Species boundaries among extremely diverse and sexually dimorphic *Arrenurus* water mites (Acariformes: Hydrachnidiae: Arrenuridae)

**DOI:** 10.1101/2021.04.04.438411

**Authors:** Mariusz Więcek, Łukasz Broda, Heather Proctor, Miroslawa Dabert, Bruce P. Smith, Jacek Dabert

**Author notes:** E-mail addresses (Łukasz Broda) (Heather Proctor), (Miroslawa Dabert), (Bruce P. Smith), (Jacek Dabert).

## Abstract

*Arrenurus* (Arrenuridae) is the most species-rich genus of mites with about 950 named species that inhabit standing, and to a lesser extent, running water habitats around the world. To date, distinguishing species of *Arrenurus* has been based on male reproductive morphology. Here, we use morphological and molecular approaches to examine species boundaries among 42 named species of *Arrenurus*, including four named species that have colour variants (red and green *A. americanus*, and red and blue *A. intermedius, A. manubriator* and *A. apetiolatus*), and two unnamed morphospecies. In this study, we examine male genital structures with the use of SEM techniques, and apply mitochondrial (COI barcode region) and nuclear (28S rRNA) gene fragments to test whether male morphology reflects species boundaries in *Arrenurus* assessed by molecular analyses. Our results reveal that male reproductive morphology parallels species boundaries as judged by molecular data. We discuss the cases of genetically poorly diversified, yet morphologically clearly defined named species. Moreover, we show that based on the species we examined, colour morphs within otherwise morphologically similar specimens represent within-species variation and, in the absence of other diagnostic traits, colour itself can be misleading in distinguishing species. Our outcomes on molecular taxonomy of *Arrenurus* provide a background for testing hypotheses about speciation rate in water mites.

## 1. Introduction

Water mites (Actinotrichida: Parasitengonina: Hydrachnidiae) are one of the most species-rich groups of arthropods in fresh water (Di Sabatino et al., 2008); however, despite being widespread and taxonomically diverse, they are poorly studied compared to freshwater insects and crustaceans (Martin et al., 2010; Proctor et al., 2015). The number of studies of water mites that incorporate molecular data is growing, and previously unrecognized diversity has been frequently revealed (e.g. Stålstedt et al., 2013; Pešić et al., 2017; Pešić & Smit, 2017; García-Jiménez et al., 2017). However, considering more than 6,000 described species found on all continents except Antarctica, placed into 57 families, 81 subfamilies and over 400 genera (Di Sabatino et al., 2008), only small number of water mite taxa have been involved in those studies.

Based on molecular clock estimates, the highly successful and cosmopolitan genus *Arrenurus* Dugès (Dugès, 1834) began to diversify 15 MYA (Dabert et al., 2016), and clearly underwent an explosive speciation that made it the most species-rich genus of arachnids that comprises approximatelly 950 named species worldwide (Cook, 1974; Smit, 2012; Gerecke et al., 2016). Depending on the species, larvae parasitize hosts from the insect orders Odonata, Diptera, and, more rarely, Coleoptera (Cook, 1974; Böttger & Martin, 2003). Deutonymphs and adults are predators of ostracods, cladocerans and to a lesser extent small insect larvae (Proctor & Prichard, 1989).

Sperm transfer behaviour in *Arrenurus* is often complex and species-specific (Proctor, 1992). It involves close pairing between males and females, and is correlated with modifications of the male’s hindbody (the ‘cauda’) and hind legs (presence or absence of a spur-like extension of the genu designed to clasp the female’s legs), and presence or absence of the petiole, a structure involved in transferring sperm (Proctor & Wilkinson, 2001). In those *Arrenurus* for which sperm transfer behaviour has been described, species whose males have well-developed petioles use them as intromittent structures to introduce sperm into the genital tract of females, whereas those that do not have well-developed petioles deposit spermatophores on the substrate and then manoeuvre the genital opening of the female overtop the sperm packet (e.g. Proctor, 1992; Proctor & Smith, 1994; Proctor & Wilkinson, 2001). Females of petiolate species may have less control over whether they take in sperm from a particular male. Because uptake of sperm seems to be more under female control in species lacking a well developed intromittent organ, female choice is supposed to be the dominant force of selection in these species, whereas sexual conflict is assumed to play a bigger role in species with males equipped with elaborate intromittent organs (Proctor & Smith, 1994; Proctor & Wilkinson, 2001). Female *Arrenurus* show relatively little variation in body shape, and species-level taxonomy of *Arrenurus* is based on the external reproductive morphology of males (Smit, 2012; Gerecke et al., 2016).

In nature, species arise in many ways including geographical isolation through to diversification of ecological niches and ending with rarely examined sexual selection (De Queiroz, 2007). The last scenario is still to a great extent a riddle, because it is especially difficult to test experimentally (Mendelson & Shaw, 2005). Theory predicts that genitalia (Eberhardt, 1985), pheromonal communication (Lassance et al., 2019) and courtship behaviour (Mendelson & Shaw, 2005) can evolve rapidly, increase sexual isolation and accelerate speciation in animals (Arnqvist et al., 2000; Janicke et al., 2018). While in some species of *Arrenurus* males differ little from females, most of the diversity in the genus composes of species with males that have extravagant dimorphism. Hence, subgenus *Arrenurus* s. str. with males characterized by hindbody and hind legs modifications and presence of the elaborated intromittent organ groups about 300 species. Further 300 species belong to the subgenus *Megaluracarus* that composes of males with exaggerated, very elongated and modified hindbody and hind legs designed to grasp and hold females during copulation. However, the least modified male phenotype, with males that does not differ very much from females is the least frequent (54 species (subgenus *Truncaturus*)); data found at website: https://bugguide.net/node/view/428959).

Here, we test species boundaries among *Arrenurus* water mites based on morphological investigation and molecular analysis of DNA barcode sequences, i.e. the mitochondrial cytochrome *c* oxidase subunit I (COI) and the hypervariable D2 region of 28S rRNA gene (28S rDNA). In order to evaluate species borders among *Arrenurus* using phenotypic characters we examined the variation of morphological secondary sexual traits in males, which is the basis for the traditional classification at the subgenus and species level (Cook, 1974). In addition, we examined what appeared to be intraspecific colour phenotypes of some of the examined species to ask whether distinctive colour forms among otherwise morphologically similar specimens might represent cryptic species.

## 2. Materials and methods

### 2.1. Water mite collecting and morphological analyses

In total, 262 *Arrenurus* mite specimens were collected in North America and Europe in various types of freshwater habitats including springs, streams, rivers, lakes, ponds, and temporary water bodies (Table 1). Most of the North American species came from water bodies located around the Queen’s University Biological Station (Ontario, Canada; 44°34’03.6”N 76°19’26.6”W), with one species collected on a private property near Elk Island National Park (Alberta, Canada; 53°39’23.7”N 112°45’37.0”W). Specimens of *Arrenurus* (*Megaluracarus*) *manubriator* Marshall that were originally from San Marcos River (Texas, US) and Lake Opinicon (Ontario) were taken from separate laboratory cultures maintained by B.P.S. in the Department of Biology, Ithaca College, New York, U.S.A. The collection sites in Europe were located in the Netherlands, Germany, Austria, Poland, and Italy (Table 1).

**Table 1.**
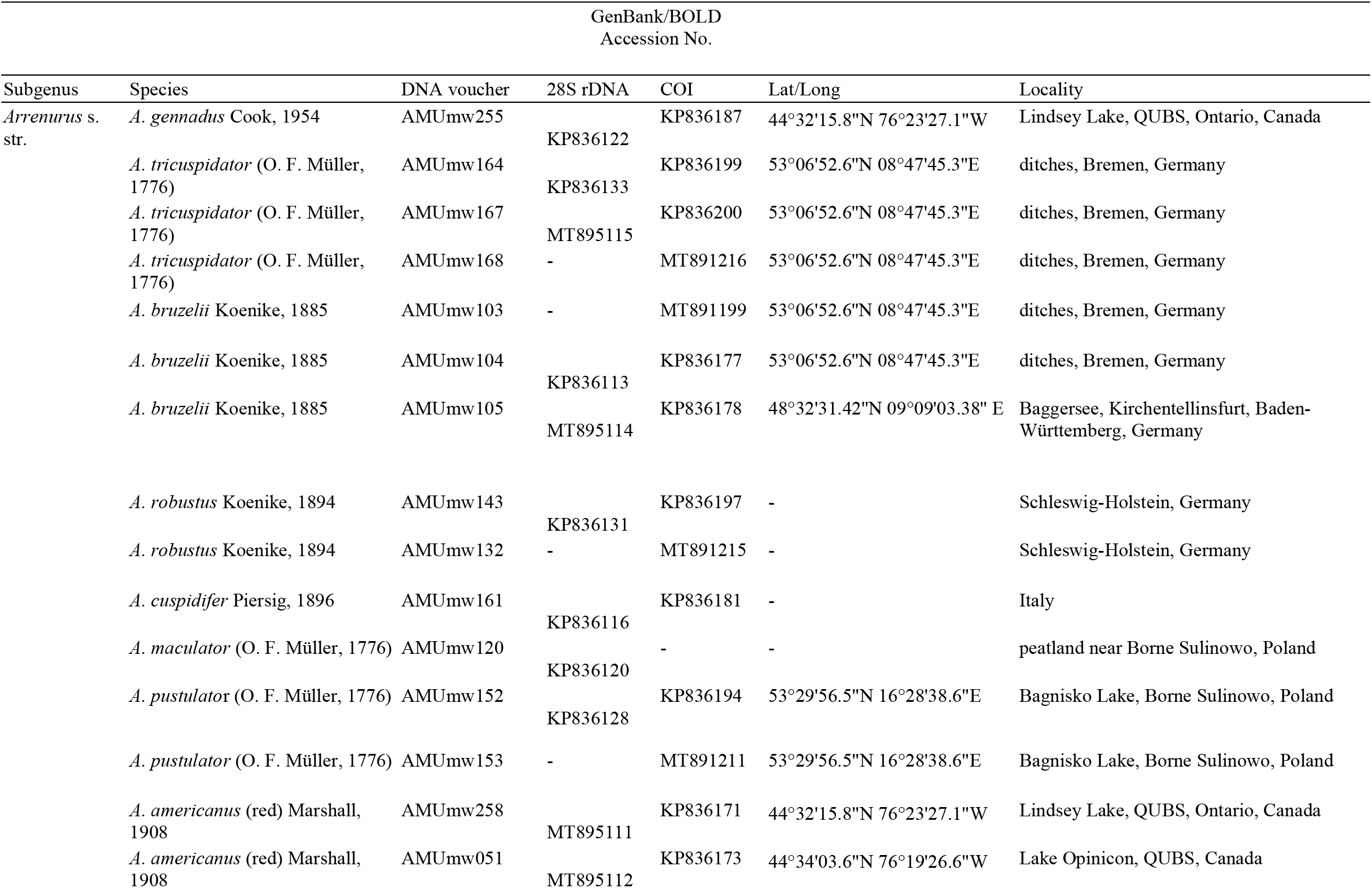
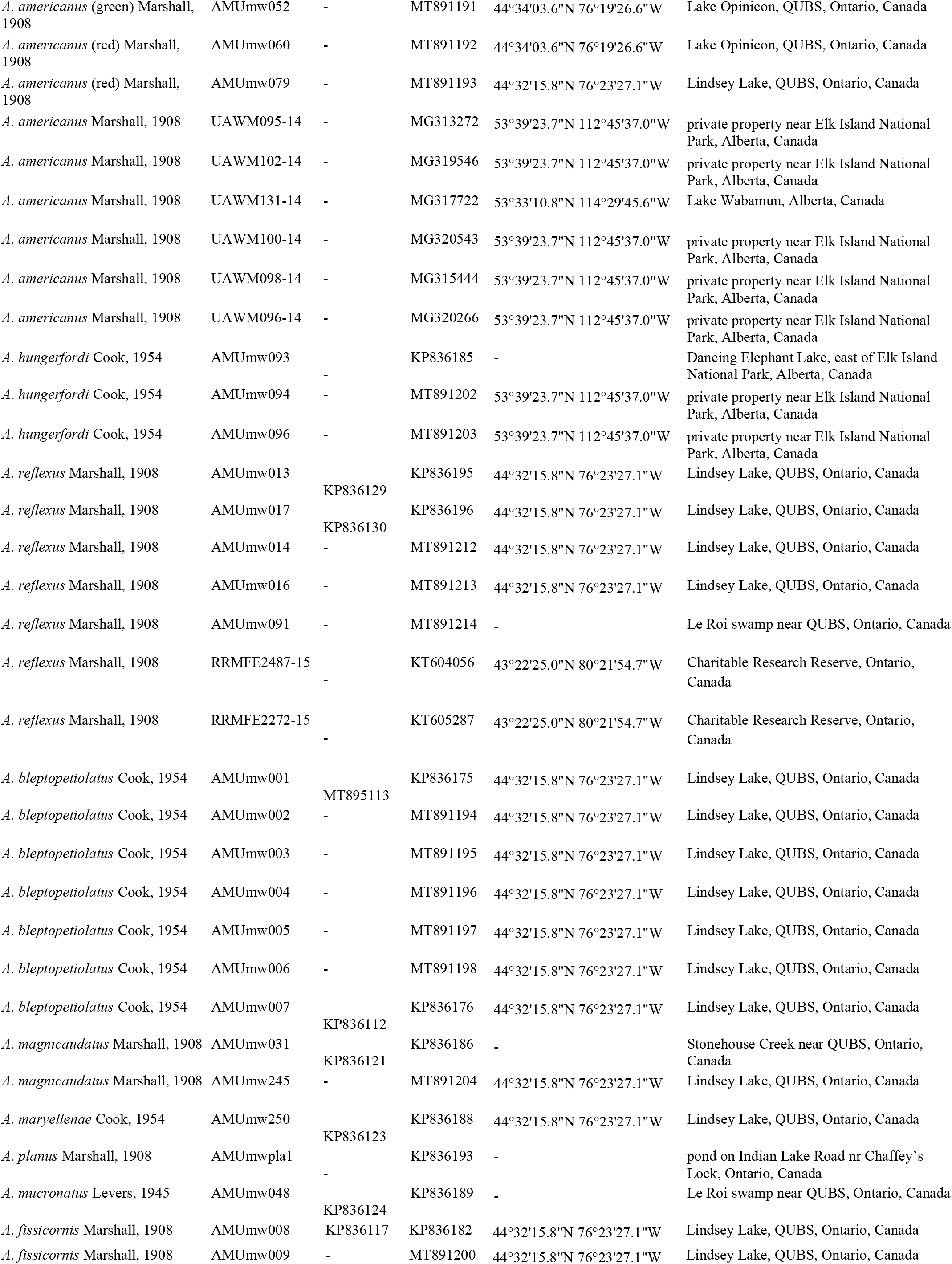
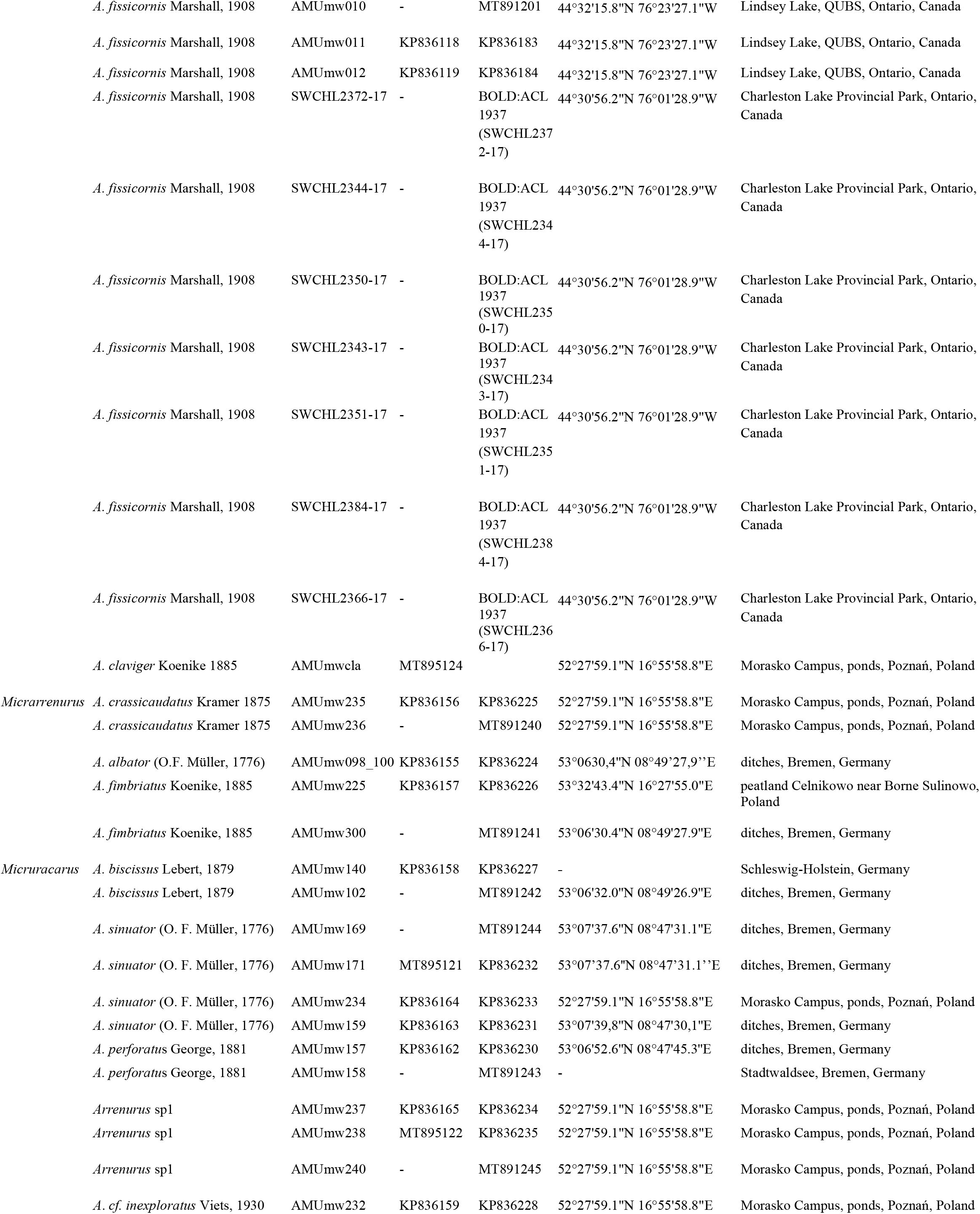
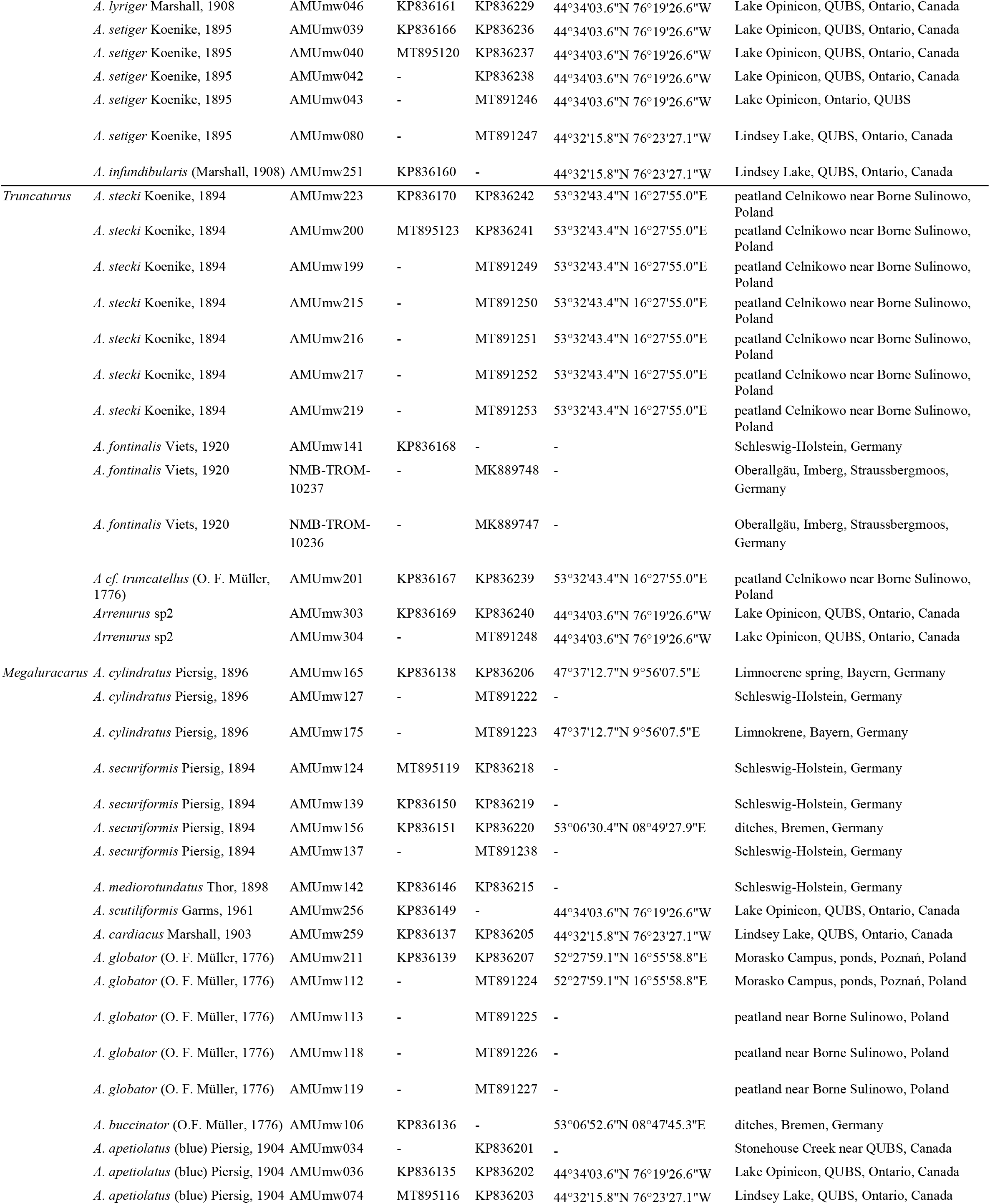
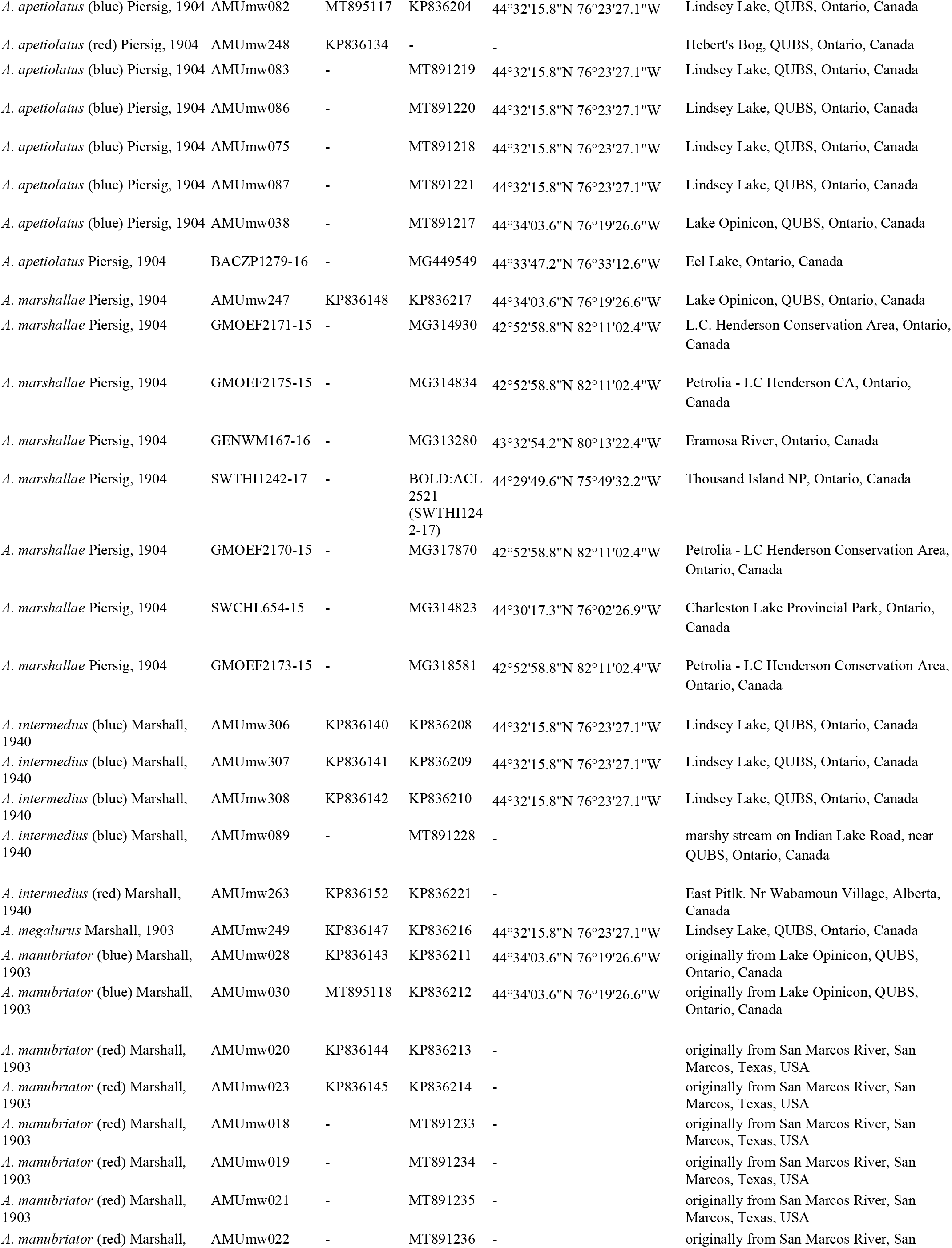
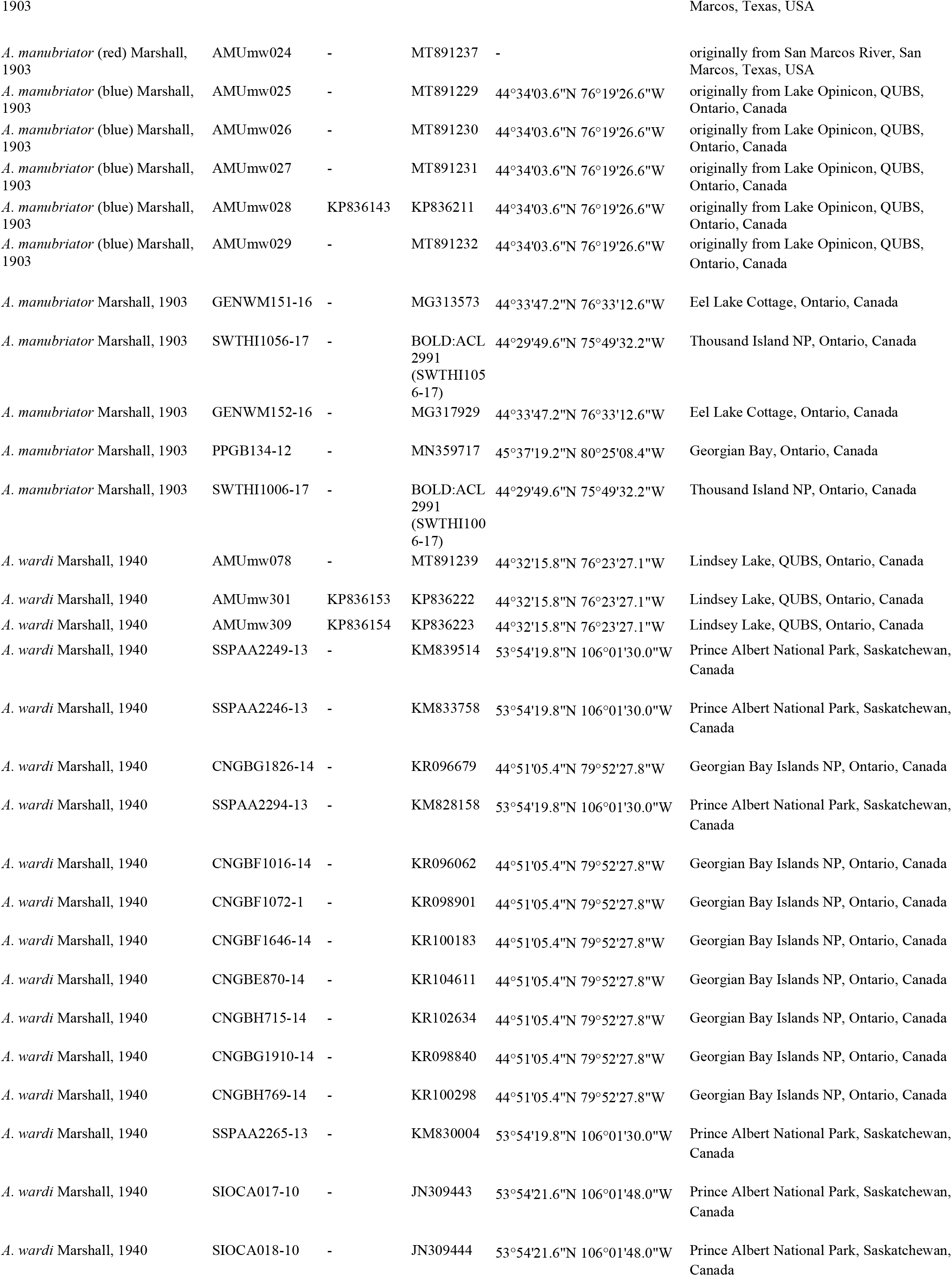
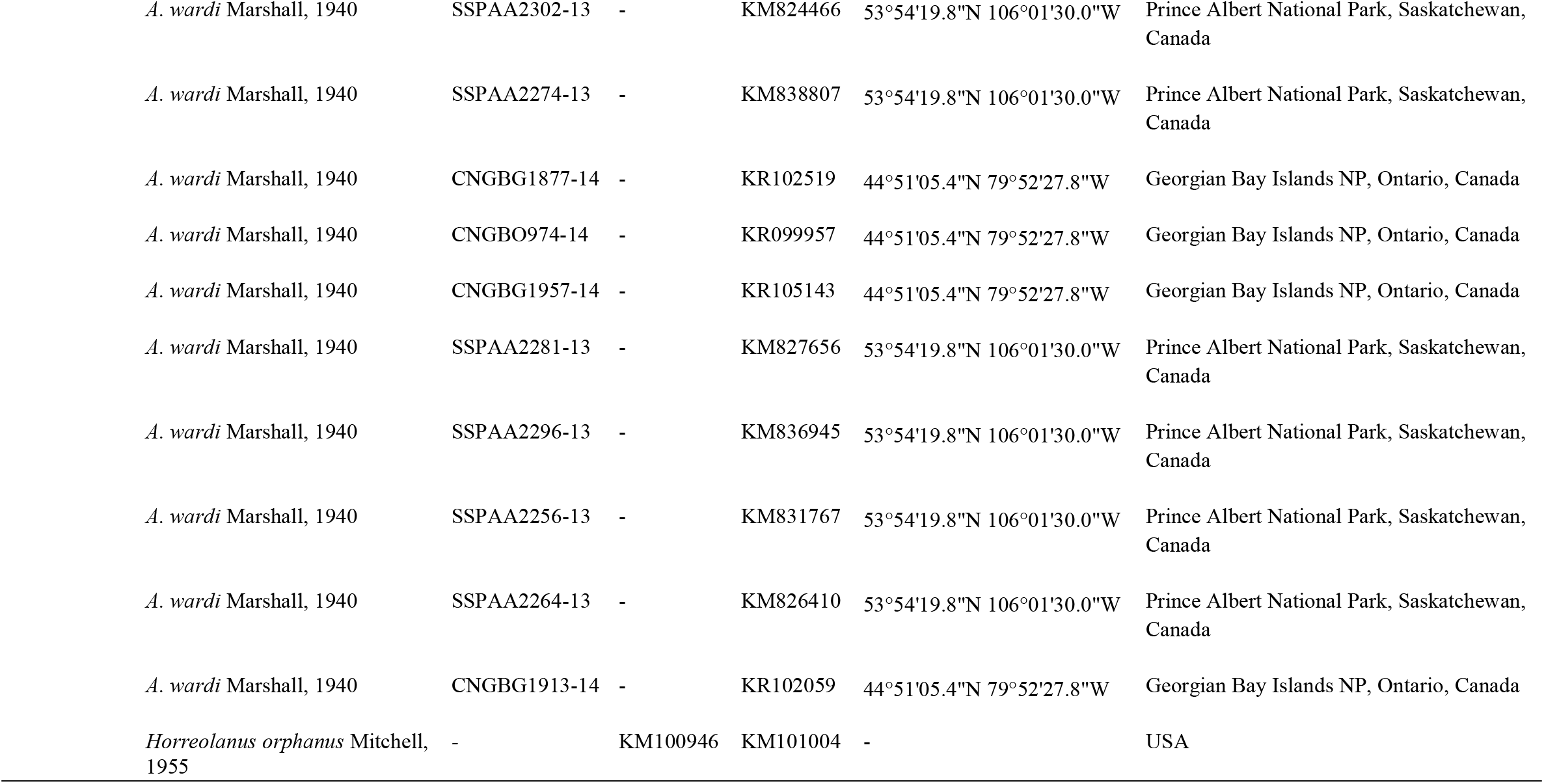
Sampling of *Arrenurus* spp. used in DNA barcoding. ‘Species’ is the *a priori* assignment that is based on morphology. Voucher information and accession numbers of sequences obtained in this study and downloaded from GenBank and BOLD are given.

Most samples were collected using a net with mesh size 250 μm, but some were collected using underwater light traps. In the laboratory, live water mites were sorted under a stereoscope microscope and preserved in 96% ethyl alcohol. European water mites were identified to species using key of Viets (1936), whereas North American ones were identified with keys by Cook (1954a, 1954b, 1955). A number of North American individuals included in the molecular analyses were determined with the assistance of Dr. Ian Smith from the Canadian National Collection of Insects, Arachnids and Nematodes in Ottawa. Only males were identified to species based on morphology; species represented only by female specimens were identified by matching COI and D2 28S rDNA sequences with those from conspecific males. When possible, representatives of named and putative species were examined using scanning electron microscopy. After dehydration through an alcohol-HMDS (hexamethyldisilazane) series, these mites were mounted on stubs using double-sided tape, sputter coated with gold, and examined using a JEOL 630 I field emission scanning electron microscope (SEM) in the Department of Earth and Atmospheric Sciences, University of Alberta. The layout of SEM images was prepared using Photoshop 6.0. Specimen and DNA vouchers from this study are deposited in the Department of Animal Morphology, Adam Mickiewicz University in Poznań, Poland.

### 2.2. DNA amplification and sequencing

Total genomic DNA was isolated from individual mites using the nondestructive method described by Dabert et al. (2008). The COI gene fragment was amplified using bcdF01 (5’-CATTTTCHACTAAYCATAARGATATTGG-3’) and bcdR04 (5’-TATAAACYTCDGGATGNCCAAAAAA-3’) primers (Dabert et al., 2010). The D2 region of the 28S rDNA was amplified with 28SF0001 (5’-ACCCVCYNAATTTAAGCATAT-3’) and 28SR0990 (5’-CCTTGGTCCGTGTTTCAAGAC-3’) primers (Mironov et al., 2012). PCR amplifications were carried out in 10 μl reaction volumes containing 5 μl Type-it Microsatellite PCR Kit (Qiagen, Hilden, Germany), 0.5 μM of each primer, and 4 μl (1-5 ng) of DNA template using a thermocycling profile of one cycle of 5 min at 95 °C followed by 35 steps of 30 sec at 95°C, 1 min at 50°C, 1 min at 72°C, with a final step of 5 min at 72°C. After amplification, the PCR reactions were diluted with 10 μl of water and 5 μl was analysed by agarose electrophoresis. Samples containing visible bands were purified with exonuclease I and FastAP Alkaline Phosphatase (Thermo Scientific) and sequenced using a BigDye version 3.1 kit and ABI Prism 3130XL Genetic Analyzer (Applied Biosystems), following the manufacturer’s instructions. Trace files were checked for accuracy and edited with ChromasPro v. 1.32 (Technelysium Pty Ltd.). The sequences generated in this study have been published in GenBank under accession numbers listed in Table 1.

### 2.3. Sequence analyses and species delimitation

From a total number of 262 *Arrenurus* specimens collected in this study, 129 were successfully sequenced with respect either to mitochondrial or nuclear marker (Table 1). The COI dataset obtained based on specimens collected in this study consisted of 123 sequences belonging to 38 named *Arrenurus* spp. (including four species possessing colour variants) and two unnamed morphospecies, and had length of 537 bp with 241 variable nps. Furthermore, we compiled a joint dataset consisting of 123 sequences obtained in this study and 54 haplotypes (from the total 196 downloaded sequences representing *Arrenurus* species) gathered from GenBank (https://www.ncbi.nlm.nih.gov/genbank/) and BOLD System (http://www.boldsystems.org/) (Table 1). The haplotype sequences were selected with the use of ALTER software (http://www.sing-group.org/ALTER/). The final alignment length and also overlapping homologous region in the joint dataset was 471 bp (221 variable nps). Since four sequence haplotypes of *A. planus* Marshall (MG312595, MG320457, MG312775, HQ924310) that were gathered from BOLD did not match conspecific sequences from this study, which could suggest possible identification errors, we excluded these sequences from the final alignment and further analyses. The D2 28S rDNA alignment comprised 68 sequences representing 41 named *Arrenurus* spp. and two morphospecies and gave 660 bp and had 241 polymorphic characters, including 29 indels. Sequences were aligned with Clustal X 2.0.10 (Larkin et al., 2007) and trimmed in GeneDoc v. 2.7.0 (Nicholas & Nicholas, 1997).

NJ tree for species-delimitation was calculated based on 177 sequences (471 bp) obtained in this study (123 sequences) and gathered from GenBank and BOLD (54 sequences) with the Kimura 2-parameter model (Kimura, 1980) in MEGA X (Kumar et al. 2018). Support for tree branches was calculated by the nonparametric bootstrap method (Felsenstein, 1985) with 1000 replicates. The critical value of bootstrap support ≥ 70% was considered to support monophyly (Douady et al., 2003). We included (Arrenuroidea Bogatiidae: *Horreolanus orphanus* Mitchell) as an outgroup species. COI (K2P) distances were reconstructed on COI alignment comprising 123 sequences belonging to specimens collected only in this study with the length of 537 nucleotide positions (nps) using MEGA X (Kumar et al., 2018). K2P distances for 68 D2 28S rDNA sequences (660 nps) from this study were computed in MEGA X (Kumar et al., 2018). The Automatic Barcode Gap Discovery (ABGD) method was applied to detect a barcode gap in the pairwise distance distribution and to sort the 123 COI sequences into hypothetical species (Puillandre et al., 2011) with the use of webbased program (https://bioinfo.mnhn.fr/abi/public/abgd/) with default settings except X (relative gap width) set to 1, because higher values failed to detect more than one group. Gene genealogies were estimated on the basis of 123 COI sequences from this project in TCS 1.21 (Clement et al., 2000) using statistical parsimony networks (SP) (Templeton et al., 1992). The 95% connection limit for species boundaries was applied in searching for putative species (Hart & Sunday, 2007). Additionally, the probability of species distinctiveness was estimated for closely related species by two measures estimating probability that the observed branching structure of the haplotypes originated due to random coalescence processes and not speciation events: reciprocal monophyly P_AB_ (Rosenberg, 2007) and Randomly Distinct P_RD_ (Rodrigo et al., 2008) as implemented in Geneious 9.1.5 species delimitation plugin (Masters et al., 2011). Editing of trees was performed using MEGA X (Kumar et al., 2018) editing tools and Inkscape 0.48.4-1 (Harrington, 2004-2005). Box plots were calculated in order to obtain medians and quartiles and to get a visual interpretation of genetic distances using the statistical software PAST 4.03 (Hammer et. Al, 2001).

## 3. Results

### 3.1. Male sexual morphology and colour variants

Examination of morphological structures, in particular structures used by males for the process of sperm transfer, allowed us to assign 129 individual mites (only specimens from which DNA barcode sequences were obtained were analysed for morphology) to 42 named species of *Arrenurus*, which included colour variants of four described species that could potentially represent unnamed species: red and blue *A. (Megaluracarus) apetiolatus* Piersig, red and green *Arrenurus (Arrenurus) americanus* Marshall, red and blue *Arrenurus (Megaluracarus) intermedius* Marshall, and red and blue *A. (Megaluracarus) manubriator*. We also found two morphospecies that did not key to known species, which we refer to as *Arrenurus* sp. 1 represented only by female specimens which was not sufficient to identifiy them at a species level, and *Arrenurus* sp. 2 consisted of female and male specimens with males possessing unmodified hindbody not clearly demarcated from the body proper and therefore resembling representatives of subgenus *Truncaturus* (Fig. 2C). In the examined species four main male phenotypes were identified: a) males with short and least modified cauda (e.g. *A. (Truncaturus) stecki* Koenike, *A. (Truncaturus) fontinalis* Viets), b) males with complex, but short cauda with medial cleft and petiole (sometimes with functionally not defined membranous structure) (e.g. *A. (Micruracarus) perforatus* George, *A. (Micruracarus) biscissus* Lebert, Fig. 2B), c) males equipped with very elongated and exaggerated hindbody without medial cleft and pygal lobes (e.g. *A. (Megaluracarus) buccinator* (Müller), *A. (Megaluracarus) globator* (Müller) (Fig. 1A, Fig. 2F), d) males that possess elaborated cauda with humps, clearly marked pygal lobes and complex petiole (subgenus *Arrenurus* e.g. *A. bleptopetiolatus* Cook, *A. magnicaudatus* Marshall, Fig. 2D, E, G). Moreover, we found species with males that deviate from above mentioned phenotypes and have short cauda with pygal lobes and membranous sub-petiolar cavity (*A. (Micrarrenurus) albator* (Müller) (Fig. 2A), *A. (Micrarrenurus) crassicaudatus* Kramer). Sexual dimorphism and subset of the diversity of male reproductive structures are presented in Figs. 1 and 2.

**Figure 1.**
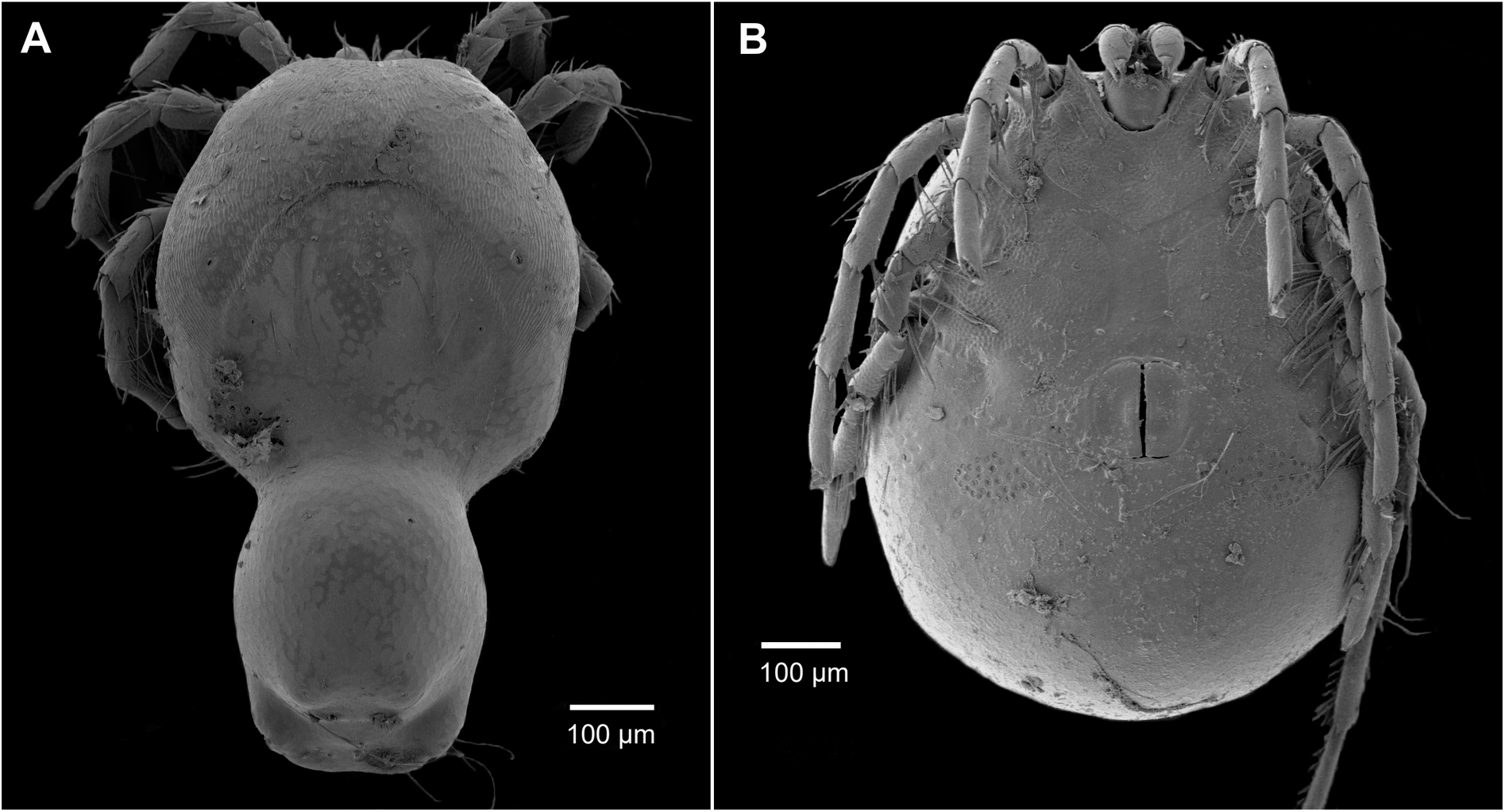
Sexual dimorphism in *Arrenurus (Megaluracarus) globator:* A. male with elongated and modified hindbody (“spur” on IVth leg not visible), B. female possesses very little diversified body and lacks cauda.

**Figure 2.**
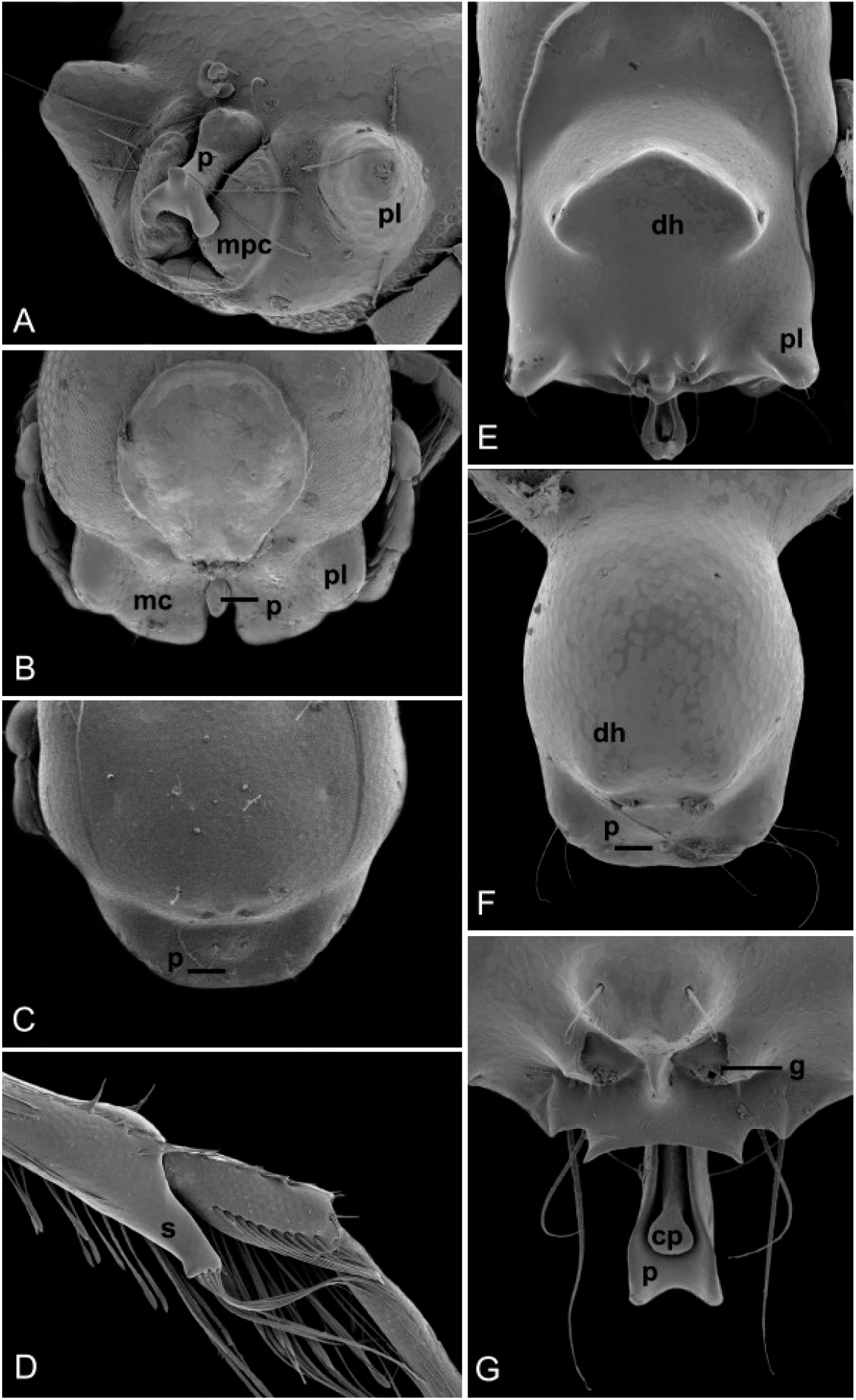
Male hindbody (cauda), intromittent, and grasping organs in *Arrenurus* spp.; A. hindbody of *A. (Micrarrenurus) albator* with membranous sub-petiolar cavity, petiole without central piece, B. cauda of *A. (Micruracarus) biscissus* equipped with small and partly membranous petiole located in elaborated medial cleft, C. slightly elongated hindbody of *Arrenurus (Truncaturus)* sp. 2 with peg-like petiole, D. IVth leg of *A. (Arrenurus) bicuspidator* with spur (clasper organ), E. elaborate cauda with pygal lobes and petiole, *A. (Arrenurus) magnicaudatus*, F. very elongated and tubular cauda with peg-like petiole of *A. (Megaluracarus) globator*, G. intromittent organ with central piece, *A. (Arrenurus) bicuspidator*; abbreviations: cp — central piece of petiole, dh — dorsal hump, g - glandularium, mc — medial cleft, mpc - membranous sub-petiolar cavity, p — petiole, pl — pygal lobe, spur (grasping structure).

### 3.2. Molecular species delimitation and morphospecies with high genetic distance

The examined 42 named species formed distinct and well supported clusters in the NJ tree inferred based on the COI seqences obtained in this study and gathered from GenBank and BOLD (Fig. 3). Levels of interspecific COI genetic differentiation based on K2P distances were in most cases well above 11% (Table S1) and had an average value of 26.06% ± 2.06% (mean ± standard error) and median value was 24.65%. The intraspecific distances ranged from 0.0% (e.g. *Arrenurus (Megaluracarus) globator)* to 2.09% (*Arrenurus (Arrenurus) bruzelii* Koenike). The interspecific D2 28S rDNA distances had an average value 14% ± 0.98 (mean ± standard error, Table S2) and median value of 13.34%. However, genetic differentiation expressed in genetic distances varied within differend subgenera (Fig. S2). The ABGD method was applied in order to estimate a barcode gap between K2P distances (Puillandre et al., 2011), which was identified between 3 and 7% (Fig. 4A, B). The analysis of 123 sequences from this study revealed 38 *Arrenurus* species in all initial partitions (Fig. S5) and 39-55 species in recursive partitions. Furthermore, in the network analysis the 123 sequences displayed 81 unique haplotypes that formed 21 distinct networks of which 17 corresponded with single named species and one unidentified morphospecies (Fig. S3), one network contained haplotypes from more than one species and two represented split networks of single species (Fig. S4). In conclusion, all applied methods confirmed species status of 35 out of 42 named species, and unnamed *Arrenurus* sp. 1, *Arrenurus* sp. 2 were identified as genetically differentiated at the species level.

**Figure 3.**
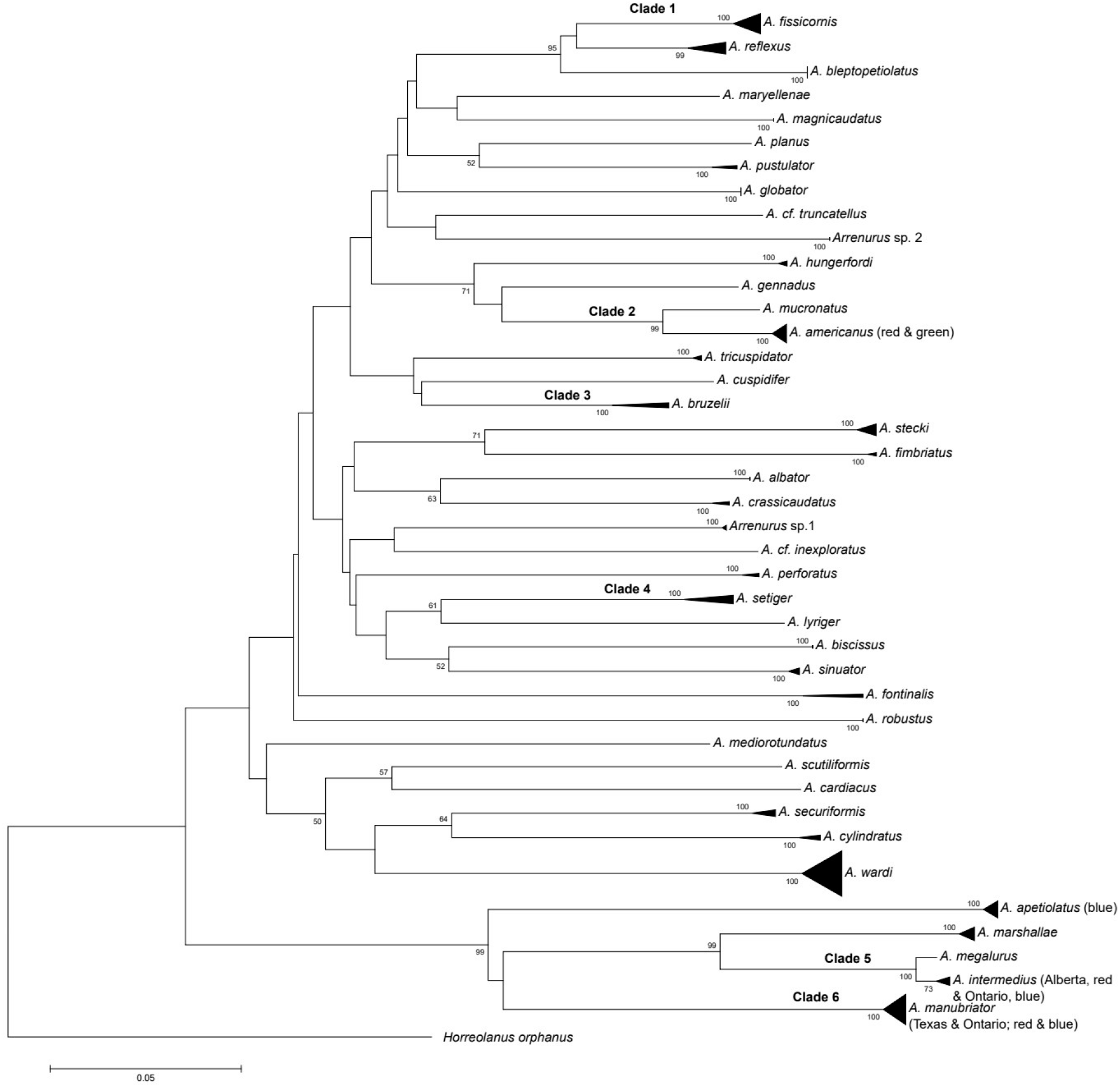
Kimura 2-parameter neighbor-joining tree of *Arrenurus* spp. based on 177 COI sequences from this study and data downloaded from GenBank and BOLD; bootstrap supports are given next to branches (only bootstrap values > 50 are shown). Clades consisted of more than one specimen are condensed; clades 1 - 6 represent closely related lineages - species delimitation analyses are summarized in Table 2. The NJ tree with expanded clades is shown in Supplementary Material, Fig. S1. *Horreolanus orphanus* is an outgroup species. See Table 1 for sequence codes and accession numbers.

**Figure 4.**
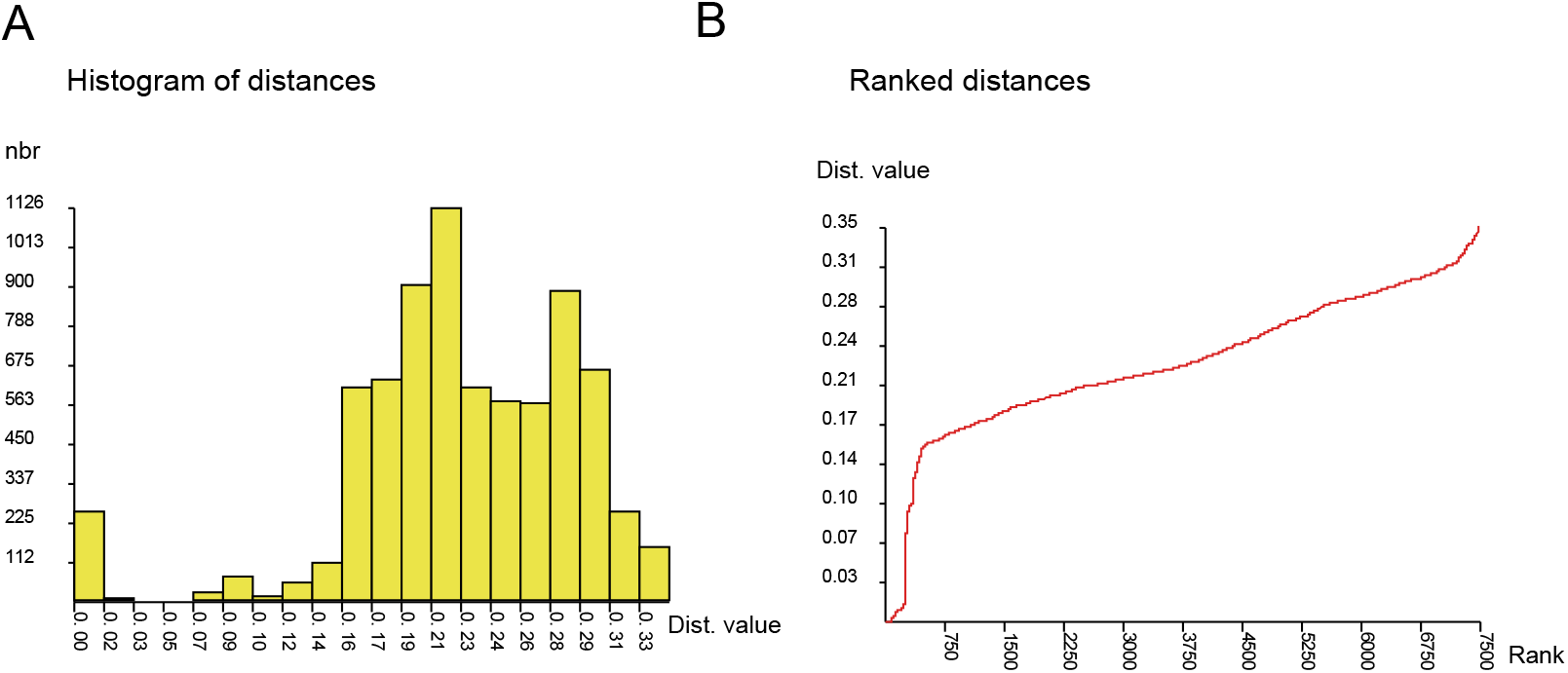
Results of ABGD species delimitation for 123 COI sequences from this study that belong to *Arrenurus* spp.; A - frequency histogram of K2P pairwise distances, B - ranked distances.

### 3.3. Morphospecies and colour variants with low genetic divergence

In the present study, except for genetically well separated species, pairs of named species characterized by low genetic distances were identified. Hence, relatively low diversification of COI sequences was found in closely related *Arrenurus (Arrenurus) mucronatus* Levers and *A. (Arrenurus) americanus* (5.1%). This species pair showed a very low level of 28S differentiation (0.15%). Similarly, low 28S distances were observed among following *Arrenurus* s.str. species pairs: *A. gennadus* Cook *-A. mucronatus* (0.00%, consistently separated as distinct species based on the COI fragment), *A. americanus - A. gennadus* (0.15%, distinct species based on the COI) and *A. maryellenae* Cook - *A. magnicaudatus* (0.31%; recognized as separate species in COI based analyses). The ABGD initial partitions did not separate *A. mucronatus* from *A. americanus* (Fig. S5). Furthermore, the red and green *A. americanus* showed intraspecific genetic variation. In the network analysis haplotypes *A. americanus* (green) and *A. americanus* (red) formed a single network, but *A. mucronatus* remained unconnected (Fig. S3). Rodrigos’s P(RD) suggested that both colour forms of *A. americanus* were one single species (Table 2).

**Table 2.** Results of species delimitation analyses based on COI DNA sequences obtained in this study. Remaining well-supported clades within species were concordantly recovered as conspecific by all methods. Abbreviations used (+): separate species, (-): the same species, (?): ambiguous result.

| Clade number | Distance comparison | | | | | $P_{(AB)}$ | Conclusion |
| --- | --- | --- | --- | --- | --- | --- | --- |
| | Distance between groups (%) | Inter /Intra-group distance ratio | ABGD | TCS | $P_{(RD)}$ | | |
| 1<br>*A. fissicornis* | 1.4 (-) | 11.1 (+) | (-) | (-) | <0.05 (+) | 0.05 (+) | Conspecific, two well-supported clades of coalescent origin yet genetically poorly diversified |
| 2<br>*A. americanus* (red and green),<br>*A. mucronatus* | 4.9-5.1 (?) | 12.5 (+) | (-) | (+) | <0.05 (+) | 0.07 (-) | Speciation or hybridization of *A. americanus* and *A. mucronatus* is not finished |
| 3<br>*A. bruzelii* | 2.7-3.0 (-) | 6.25 (-) | (-) | (+) | NA | 0.33 (-) | Conspecific, well genetically separated clades of coalescent origin |
| 4<br>*A. setiger* | 2.4-2.8 (-) | 7.69 (-) | (-) | (+) | NA | 0.1 (-) | Conspecific, well genetically separated clades of coalescent origin |
| 5<br>*A. megalurus*,<br>*A. intermedius* (red and blue) | 1.3-1.7 (-) | 2.38 (-) | (-) | (-) | NA | 0.07 (-) | *A. megalurus* and *A. intermedius* are conspecific |
| 6<br>*A. manubriator* Ontario (blue) and Texas (red) | 2.18 (-) | 9 (+) | (-) | (-) | <0.05 (+) | <<0.001 (+) | Conspecific, but well separated groups being formed by speciation mechanism rather than coalescence |

In the species pair *Arrenurus (Megaluracarus) megalurus* Marshall and *A. (Megaluracarus) intermedius* (red and blue) COI distances ranged from 0.95% - 1.58% (Table S1). The ABGD initial partitions did not separate *A. megalurus* from *A. intermedius* (both red and blue) (Fig. S5), and statistical parsimony networks grouped together haplotypes of *A. megalurus* and *A. intermedius* (Fig. S4). Similarly, Rosenberg’s P_AB_ suggested that these species are conspecific (Table 2). In addition, a low 28S distance value occurred between *A. intermedius* and *A. megalurus* (0.46%). Moreover, the blue and red *A. (Megaluracarus) apetiolatus* were separated by distance of 0.31% (28S), which could suggest that two separate species are present. However, the result remains inconclusive, because COI sequences were not successfully obtained for this species. In the colour-polymorphic *A. (Megaluracarus) manubriator* red specimens originated from Texas and blue ones from Ontario were separated by 0.99% (COI) and 2.18% (28S). Rosenberg’s P_AB_ and Rodrigos’s P(RD) indicated that red and blue *A. manubriator* should be considered as separate species (Table 2). Nevertheless, the contradictory results were obtained after inclusion in the analysis sequences gathered from BOLD (Fig. S1). Results of species delimitation in species that were not concordantly recovered as conspecific are summarized in Table 2.

## 4. Discussion

Our data show that male reproductive morphology, which provides the rationale for most taxonomic decisions in *Arrenurus* tends to parallel species boundaries as judged by molecular data. The ABGD analysis identified barcode gap between 3-7%, which is lower than threshold obtained for spring-dwelling water mites by Blattner et al. (2019) (6-9%). Interspecific COI barcode distances have been estimated for congeneric vs. congeneric water mite species most frequently for > 10% (Pešić & Smit, 2017; Pešić et al., 2017; Więcek et al., 2020), and correspond to the values obtained in this study. However, noteworthy, among closely related *Atractides* species distances had values of about 5% (Pešić et al., 2020). García-Jiménez et al. (2017) obtained a statistical support for separating endemic island *Lebertia* water mites that exhibited distances of about 2% and shared the common ancestor about 4.6–5.2 Mya. The simulation study found bias against discovering young species in taxa undergoing adaptive radiation and demonstrated that single-gene thresholds can consistently discover new species with error rates of <10% if isolation was >4 million generations ago (Hickerson et al., 2006). Therefore, we suppose that low sequence divergence in species groups examined in this study may indicate young, but reproductively isolated species. In addition, the mean distance value for 28S sequences obtained in this study (14%) is in agreement with sequence diversification observed for a broad set of water mites inhabiting springs (15%K2P ± SD: 0.10%) found by Blattner et al. (2019), and in most cases paralleled results obtained with mitochondrial sequences. Interestingly, we observed low differentiation at nuclear loci among members of subgenus *Arrenurus* s. str. as compared to representatives of other examined subgenera. The subgenus *Arrenurus* str. is considered as monophyletic (when excluding members of the subgenus *Micrarrenurus* proposed later by Cassagne-Méjean, 1966) and molecular dating analysis suggested its recent origin in relation to other examined *Arrenurus* taxa (5–10 Mya Dabert et al., 2016). Similar low distance values were also obtained for young eriophyoid mite species (Skoracka & Dabert 2010). To date, there are very few studies targeting certain aspects of taxonomical assesments of *Arrenurus*, where the validity of morphospecies is tested with the application of molecular markers (e.g. Blattner et al., 2019; Alarcón-Elbal et al., 2020). However, in the recent published article of Alarcón-Elbal et al. (2020) phylogenetic relationships among *Arrenurus* species were missinterpreted. Whereas the authors stated that subgenera *Arrenurus, Megaluracarus* and *Micruracarus* are “natural” and only subgenus *Truncaturus* is an arbitrary assemblage of species, in fact non of the examined by the authors subgenera is monophyletic when interpreting results of the phylogenetic tree presented by the authors (Alarcón-Elbal et al., 2020).

We obtained equivocal results for closely related *A. (Megaluracarus) intermedius* and *A. (Megaluracarus) megalurus*, which show very low genetic differentiation. Moreover, other than colour, there is no morphological evidence for separating the blue and red *A. intermedius*. It is noteworthy that *A. intermedius, A. megalurus* and *A. (Megaluracarus) marshallae* Piersig have been considered in literature as closely related, morphologically similar species that exhibit subtle differences in male structures associated with hindbody and that tend to occur in the same habitats at the same time (Cook, 1954b; Mitchell, 1964). Furthermore, the two colour forms of *A. (Megaluracarus) apetiolatus* are probably recently-diverged species, however we have insufficient evidence given that we were unable to obtain mitochondrial sequences. Moreover, we obtained ambiguous evidence for distinguishing species in closely related *A. (Arrenurus) mucronatus* and *A.(Arrenurus) americanus* (both red and green individuals). *Arrenurus americanus* is highly variable in colour: they are predominantly either dark green or brick red, but various other colours also occur (grey, tan, etc.) with more or less a continuum of variation (B.P.S., pers. obs.). In this study, we observed intraspecific sequence differences between both colour variants of *A. americanus*. However, we found initial stages of sequence diversification between *A. americanus* and *A.mucronatus*, which may indicate the presence of a young and potentially hybridising species. Given that *A. mucronatus* has consistent differences relating to size differences and structure of hindbody and intromittent organ when compared to *A. americanus*, the low differentiation in barcode sequences could indicate that morphology evolves more rapidly than mitochondrial sequences, as would be expected under continuous directional sexual selection (Wojcieszek & Simmons, 2011). Nevertheless, other forces as stabilizing natural selection could be potentially responsible for divergence of male genitalia. In view of the fact, that male genitalia in males in *Arrenurus* are highly complex and include presumably functionally different components, it is likely that different sections of male genitalia in *Arrenurus* may be subject of different evolutionary processes as it was suggested for grasshopper species (Song & Wenzel, 2008).

We observed a clear pattern within the geographically widespread and color-polymorphic *A. (Megaluracarus) manubriator*, from which we had representatives from distinct habitats and distant regions represented by laboratory colonies established from two populations located approx. 2,500 km apart (red mites from a river in Texas vs. blue mites from a lake in Ontario). The genetic differentiation of these populations is probably the result of processes associated with speciation, as random coalescence was rejected as explanation of this divergence. However, only limited conclusions can be drawn with regard to *A. manubriator*, because split into two distinct clades observed based on specimens from this study is not retained after including sequences of specimens collected in a wider range of habitats (data from BOLD databasis). In the apparently closely related *Arrenurus* s.str. species group *A. fissicornis* Marshall - *A. reflexus* Marshall - *A. bleptopetiolatus* all applied methods were consistent and confirmed genetic separation at the species-level. Interestingly, we found well supported clade structure of coalescent origin within *A. fissicornis*, which was also present after including data from BOLD. Overall, the comparison of our data with sequences deposited in BOLD databasis was, however, limited to a few species, because only a small part of sequences was publically available.

Its clear from our study that colour is a questionable character to use when separating species: sometimes it is informative, other times it is a false lead. There are species that are highly variable, some are remarkably consistent and colour is a useful cue for identification (esp. within site). Furthermore, we observed that body colour can be either consistent within population but varying among populations (e.g., *A. intermedius*), or in other cases also highly variable within population (e.g., *A. americanus*). The occurrence of colour variants that showed within-species sequence divergence was also found in melon aphids (Lokeshwari et. al., 2014), pabble crabs (Prakash & Kumar, 2020) and sea cucumbers (Jo et. al., 2016). However, Soto-Adames (2002) revealed that most populations of springtails differing only in color pattern showed signifficant genetic divergence and thus were recognized as distinct species. We suggest that in the absence of other morphological differences, body colour itself is not a good diagnostic trait for species separation in the genus *Arrenurus*. However, in a few cases it may be a clue that there is underlying genetic differentiation (*A. manubriator*, laboratory colonies, B.P.S., pers. obs.). Although it’s possible that variation in color in some cases may be caused by local water chemistry, body colour may have adaptive value for instance as photoprotectants (red and orange carotenoid pigments) (Proctor & Garga, 2002). Although our outcomes suggest that certain pairs of named species could be conspecific, we believe that the patterns found in this study should be further tested on larger numbers of individuals from broader geographic ranges.

## Acknowledgements

The authors thank Dr. Ian Smith, Ottawa, Canada, for assistance in identification of specimens collected in Canada. We are grateful to Dr. Reinhard Gerecke, Tübingen, Germany, and Dr. Harry Smit, Leiden, The Netherlands, for the loan of many specimens. Dr. Peter Martin, Kiel, Germany, supported the study through providing specimens and discussing parasitic associations in water mites. The study is part of the International PhD Programme ‘From genome to phenotype: A multidisciplinary approach to functional genomics’ (MPD/2010/3) funded by the Foundation for Polish Science (FNP). Additional funding was provided by a Natural Sciences and Engineering Research Council of Canada (NSERC) Discovery Grant to HCP.

## Supplementary material

Figure S1. Kimura 2-parameter neighbor-joining tree of *Arrenurus* spp. based on 177 COI sequences from this study and data downloaded from GenBank and BOLD (bootstrap supports next to branches, only values > 50 are shown). There are included GenBank sequences of *Arrenurus fontinalis* since we have failed to obtain COI sequences from our samples. *Horreolanus orphanus* is an outgroup species. Clades 1 - 6 are closely related lineages and relate to results of species delimitation analyses shown in Table 2. Table 1 includes sequence codes and accession numbers.

Figure S2. The box plots show medians, quartiles and standard errors of genetic distances within the genus *Arrenurus* and within subgenera *Arrenurus* s.str., *Megaluracarus, Micruracarus, Truncaturus* and *Micrarrenurus*; species with exceptionally low distance values and colour variants were excluded from statistical analysis; A. COI (K2P) distances - *A. intermedius* vs. *A. megalurus* were excluded from statistical analysis; B. 28S (K2P) distances - *A. intermedius* vs. *A. megalurus* and *A. gennadus* vs. *A. mucronatus* were not included; subgenera *Truncaturus* and *Micrarrenurus* were represented only by 6 (3 for COI) and 3 observations, respectively.

Figure S3. Haplotype networks of the COI gene under the 95% parsimony criterion. Networks corresponding with named species and one unnamed morphospecies are shown. The size of ovals reflects haplotype frequencies. Dots on lines indicate one missing unsampled haplotype.

Figure S4. Haplotype networks of the COI gene under the 95% parsimony criterion. Networks consisted of several named species or splitted networks are presented. The size of ovals reflects haplotype frequencies. Each bar indicates one missing unsampled haplotype.

Figure S5. ABGD output tree based on 123 COI sequences of *Arrenurus* spp. from this study. The number of groups in initial partitions is presented. See Table 1 for sequence codes and accession numbers.

Table S1. Kimura 2-parameter (K2P) distances (and standard errors) for COI sequences obtained in this study calculated within (in grey) and between species.

Table S2. Kimura 2-parameter (K2P) distances (and standard errors) for 28S sequences obtained in this study calculated within (in grey) and between species.

