## Supplementary figures and images for "Species boundaries among extremely diverse and sexually dimorphic *Arrenurus* water mites (Acariformes: Hydrachnidiae: Arrenuridae)"

### Figure S1

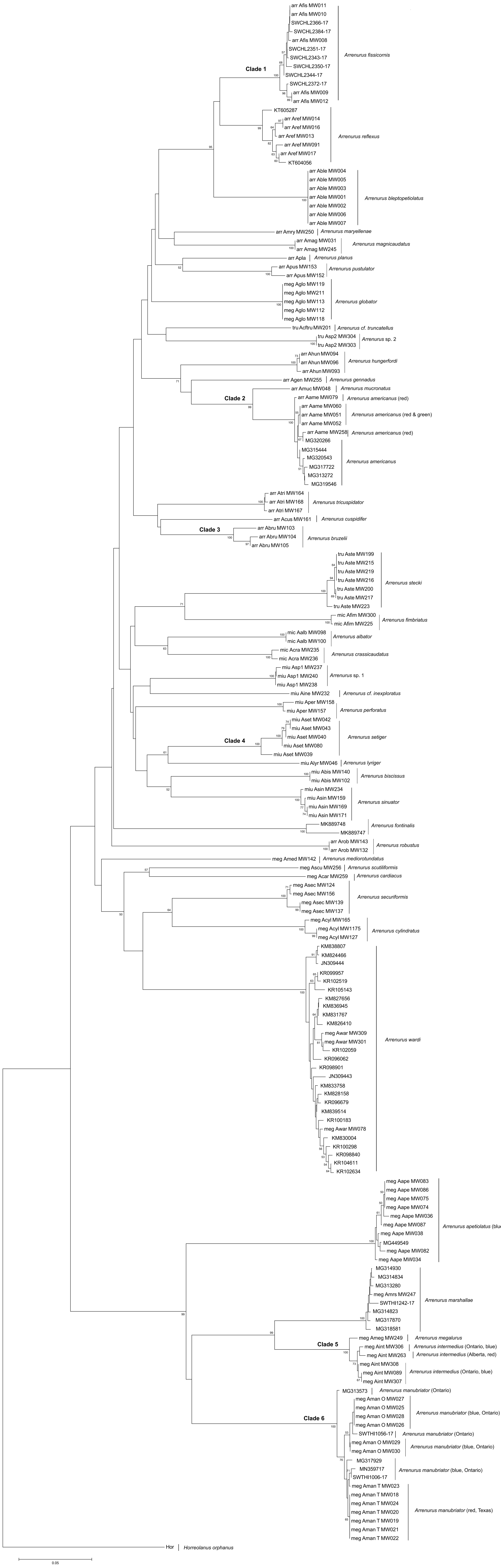

### Figure S2

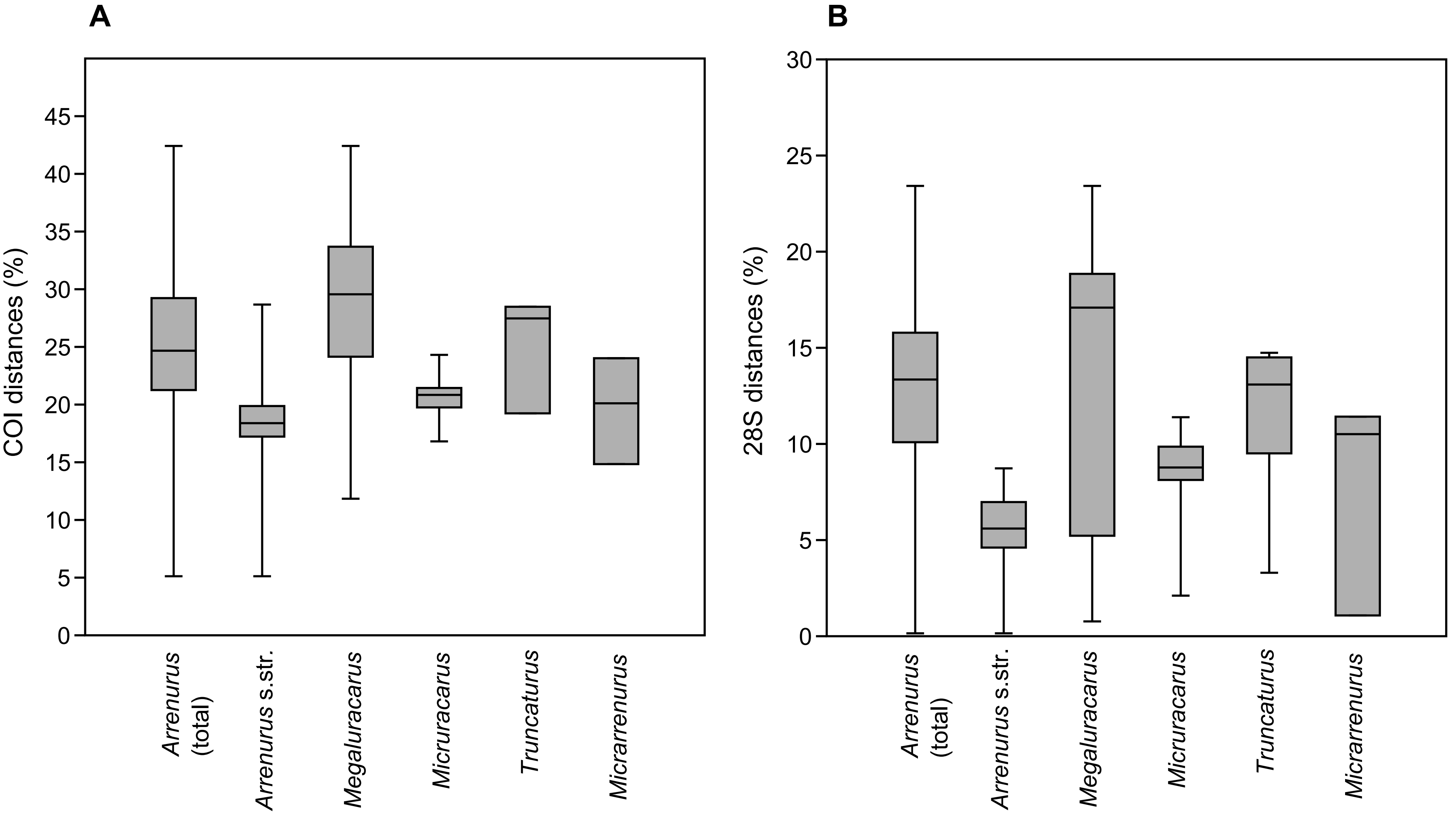

### Figure S3

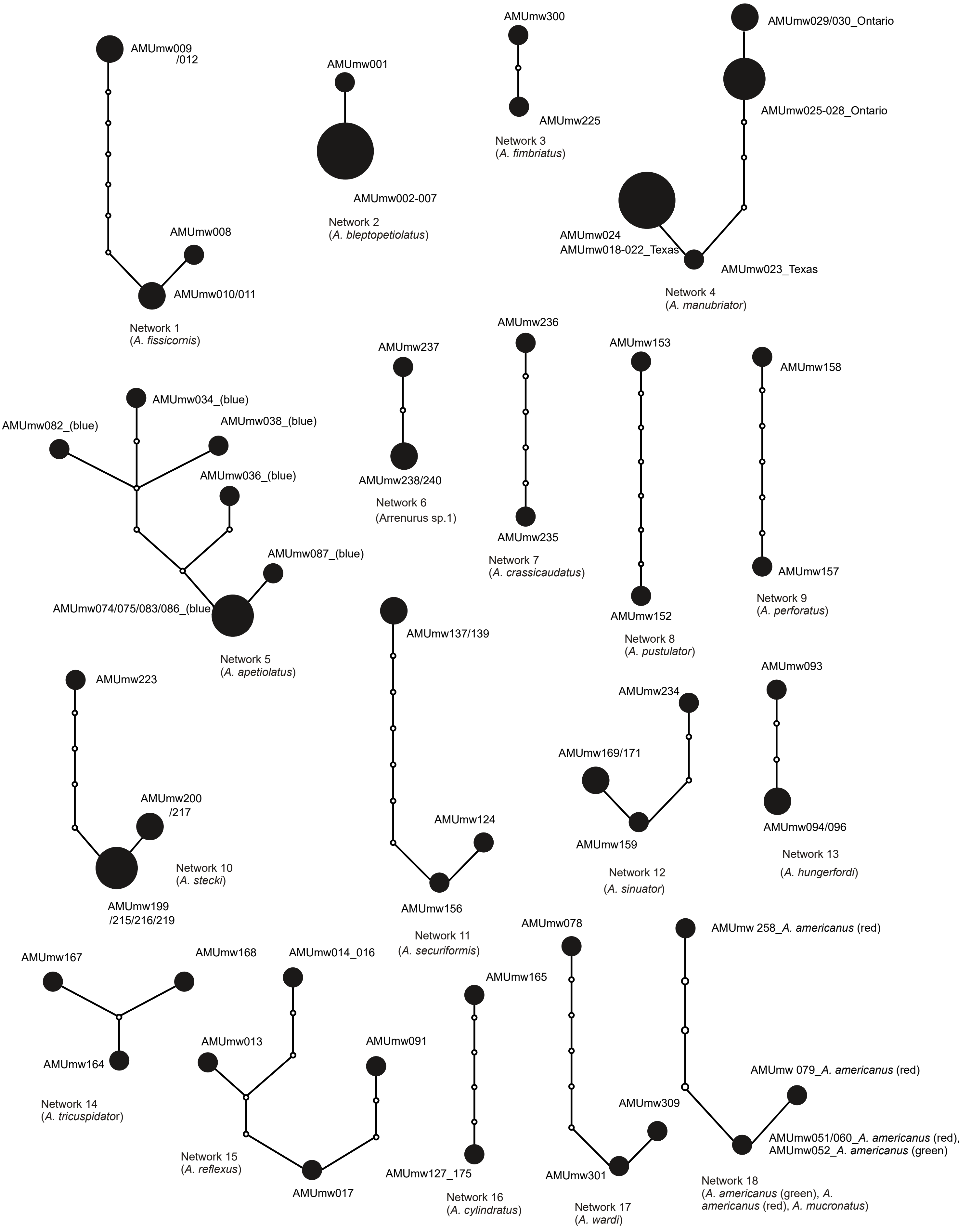

### Figure S4

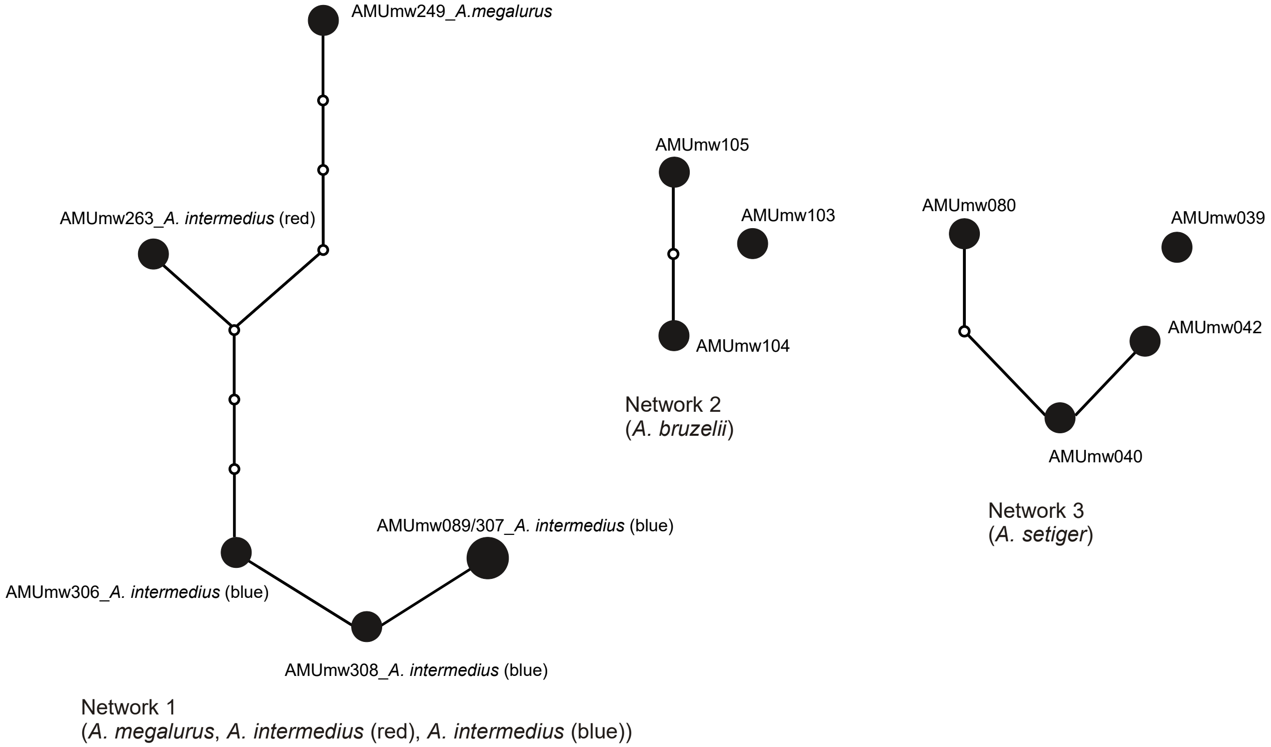

### Figure S5

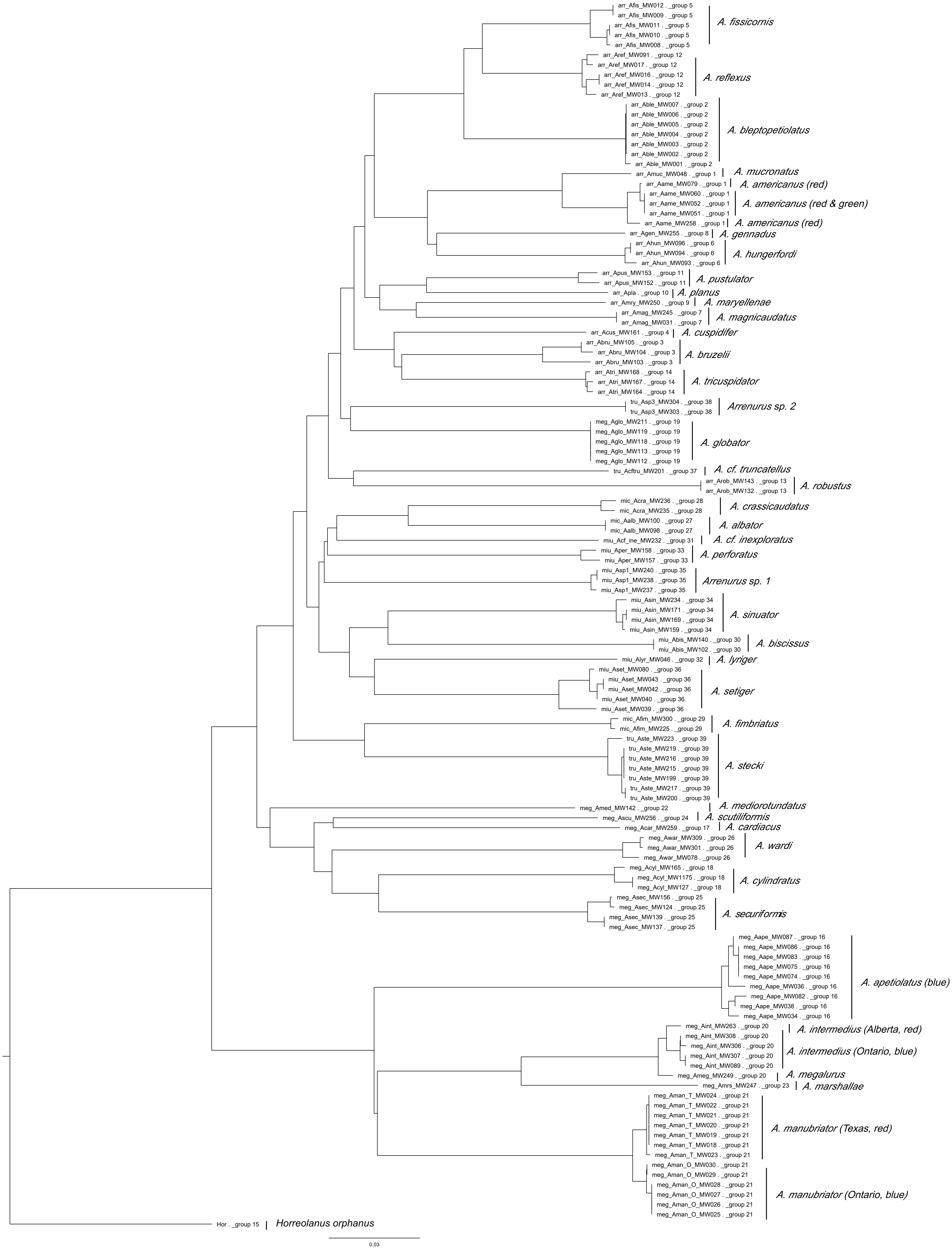
